# Ageing in house sparrows is insensitive to environmental effects

**DOI:** 10.1101/598284

**Authors:** Mirre J. P. Simons, Isabel Winney, Antje Girndt, Mark Rees, Shinichi Nakagawa, Julia Schroeder, Terry Burke

## Abstract

Variation in individual life histories, and physiology, determines the rates at which new life is generated (reproduction) and lost (death) in a population. Studying the demography of deaths thus reveals fundamental aspects of the biology of individuals within a population. We studied mortality senescence – the increase in mortality rate with age – in wild and captive house sparrows (*Passer domesticus*), and demonstrate highly similar mortality senescence in both, but markedly lower vulnerability to death (frailty) in captivity. This suggests that house sparrows have a species-specific rate of ageing that is insensitive to environmental effects. Unexpectedly, juvenile and adult mortality co-varied positively across years in the wild, indicating that mortality is not strongly density-dependent. Mortality also varied widely among years, suggesting a strong environmental effect, and we explain the observed patterns using temperature data and predation by birds of prey. We discuss how stochastic environmental effects can affect the evolution of ageing.

## Introduction

Demography can reveal unique aspects of the biology of the individuals within a population^1–5^. An interest in the demography of wild populations has recently been ignited^6–8^, driven by the question of whether wild animals show senescence. Historically, senescence was presumed to be minimal in the wild, swamped by extrinsic mortality (e.g. predation, disease, accidents, harsh weather) such that very few individuals in the wild would live long enough to show senescence^8,9^. This hypothesis has now been falsified by studies in a multitude of wild populations in which evidence has been found for both reproductive and mortality senescence^8,10,11^. In many of these studies the confounding effect of an unknown rate of dispersal with age is an important limitation, with dispersal being mistaken for mortality. The number of studies of age-specific survival remains, however, severely limited, especially compared to the many on reproductive senescence. Studying both factors together can provide a unique contribution to understanding the evolution of senescence in a comparative context^7,8^. Recent comparisons of mortality trajectories across species have revealed a wide range of patterns, from negligible senescence to a rapid acceleration in mortality with age, and this variation currently remains largely unexplained^12^.

Variation in mortality trajectories among species can result from differences in physiology and also from environmental effects and their interactions with physiology^13^. The demography of death records reveals two main components: the increasing risk of death with age – the ageing rate – and frailty, the vulnerability to death from ageing-related causes^14–16^. These parameters have both genetic and environmental components. Studies on insects, utilizing the advantage of obtaining many related offspring within a lineage, have reported heritability for both these parameters^1,17^. Another approach is to estimate the scope for environmental effects by comparing different populations of the same species or comparing mortality in the wild with that in captivity. Baboons (*Papio hamadryas*) in two different wild populations and a captive population have been shown to have different levels of frailty, yet to show a highly similar rate of ageing^18^. Similarly, the recent gain in human lifespan^19^ is due to a reduction in frailty, rather than a change in ageing rate^20^. Together, these studies suggest that there are both environmental and genetic determinants of frailty. Environmental effects on ageing rate are less common, although diet has been shown to modulate ageing rate in rodents^16^.

The underlying physiology of differential adult mortality trajectories, and especially the magnitude of the latter compared to juvenile mortality, results from selection pressures shaped by life-history trade-offs. Adult and offspring mortality risks are expected to be traded-off with the costs of reproduction, determining, for instance, the optimal clutch size and effort that should be invested in provisioning^21^. Interestingly, separating juvenile and adult survival is not usually possible because of dispersal in non-closed populations^22,23^. This limits our ability to understand the connection between demography and life-history trade-offs, and also to detect and quantify the density dependence of population dynamics^22^. For example, it would be difficult in a non-closed population to separate an effect of density on survival from an effect on density-dependent dispersal.

Here, we study the demography of an exceptionally well-monitored, closed island population^24,25^, and also of a captive population, of house sparrows^26^ (*Passer domesticus*) to (i) compare mortality senescence in the wild and in captivity. We also (ii) assess the effect of the environment on adult mortality and juvenile recruitment in the wild, factors that we were able to separate reliably due to the near-perfect resighting rate, comprehensive coverage of breeding attempts on the island, and negligible rates of immigration and emigration^24,27^.

## Methods

### Study populations

Monitoring of the house sparrows on Lundy Island (51.10° N, 4.40 W°), England, began in 1990 and has been undertaken systematically since 2000; data are presented here for the period 2000–2012. Every year, breeding behaviour in and outside nestboxes is monitored and birds are trapped using mistnets during each winter (Nov–Feb). Captures include those by researchers focusing on sparrows specifically, and those caught by the Lundy Field Society during bird surveys. Individuals are ringed before fledging with individual metal rings and a unique combination of colour rings, allowing sightings without actual captures, and individuals are sampled for blood as chicks and adults for genetic parentage analyses^10,28^. To estimate the population size and resighting probability in our study population we used actual catches^25^, *ad libitum* sightings during the breeding season and winter, structured sightings from social parentage assignment of broods using video recording^27^ and genetic sightings (genetic parentage assignment to offspring, except for the last year included, 2012, for which pedigree information was not yet available at the time of analysis). We assessed juvenile survival for all nestlings that received a metal British Trust for Ornithology ring, which they were given at an age of *ca* 12 days to focus on post-fledging survival, and we disregarded earlier deaths in the nest. Lundy is a small island (< 5.0 km long and 0.7 km wide) and is 19 km from the closest mainland shoreline, which limits dispersal from and to the island to almost non-existent^24^. The habitat on Lundy consists of a small village and farm surrounded mostly by grassland and cliffs, but with a small wooded valley. The sparrows are almost exclusively restricted to the village and the adjacent wooded valley; an excess of nestboxes was available throughout the study area. Predators consisted of occasional birds of prey that pass through during migration or, occasionally, become resident on the island in winter (see below).

The captive population of sparrows was maintained at the Max Planck Institute for Ornithology (Seewiesen, Germany) from 2005 (results include data on mortality up to 2014). Individuals were originally captured from the wild in rural Bavaria, Germany and subsequently maintained in aviaries. A proportion of the offspring born in captivity were maintained in the population and inbreeding was avoided as much as possible by transferring birds among aviaries. All birds were individually ringed and mortality was monitored daily. The specific husbandry of the birds and previous research has been described elsewhere^26,29^. The captive dataset consisted of 304 adult birds, including 170 individuals that were still alive and were therefore right-censored.

### Mortality trajectory and resighting probability

We used Bayesian Survival Trajectory Analysis (BaSTA, 1.9.2) implemented in R^30,31^. BaSTA uses a Monte Carlo Markov Chain algorithm combining Metropolis sampling for survival parameters and latent states (when times of birth or death are unknown) in a mark–recapture framework. Mark–recapture models use the missed observations of individuals known to be alive at the point of missed observation (i.e. they are observed later) to estimate the probability that an individual is sighted in the population. BaSTA combines such mark–recapture probability modelling with fitting the mortality/survival trajectory.

We fitted Gompertz and Logistic models with a bathtub shape^30,32^. Gompertz and Logistic models differ, in that Logistic models allow the mortality rate to plateau at advanced ages, whereas under the Gompertz law mortality rate continuously accelerates exponentially with age^30,32^. An increase in mortality rate with age is evidence for senescence. A bathtub shape (declining Gompertz, see equation 1) allowed early mortality, from the nestling state to adulthood, to be modelled. We selected the best model based on the deviance information criterion (DIC, which behaves similarly to AIC), and checked convergence by running each chain eight times, with each individual chain run for 1,000,000 iterations, with a burnin of 2,000 and thinning interval of 2,000. Autocorrelations of the chain were below 0.045 for all parameters in all models run. We included sex as a categorical covariate (allowing mortality parameters to vary between the sexes) in the models, because sex differences in longevity are prevalent across the animal kingdom^33^. The inclusion of this covariate might therefore improve the fit of the BaSTA model and thus the estimation of the re-sighting probability and parametric survival models. The significance of the inclusion of sex as a covariate was judged using the Kullback–Leibner divergence calibration^11,30^, which ranges from 0.5 to 1, with 0.5 indicating identical posterior distributions, and hence no effect of the covariate. We included known birth years (i.e. observed as a chick), where possible, for 2,297 of the 2,514 individuals included, for which we had 1,750 re-sightings available in 2000–2012. Known death years were included from recoveries made in the field (for 155 individuals). Sighting years were coded from 1 March until 29 February in the following year in order to include sightings up to the start of the next breeding season. Mortality trajectories in the captive population were fitted using the package ‘flexsurv’ in R in a maximum-likelihood framework^34^. Individuals that died as a result of accidents in the population, and those still alive, were right-censored. We only fitted adult mortality, because data for juvenile (under one year old) mortality were not complete, because this was not always recorded in the required detail. The parametric models fitted were limited to a Gompertz without any covariates, because in the smaller captive dataset the Logistic model did not converge and mortality deceleration was not evident in the raw data. For a direct comparison with the wild population, a simple Gompertz without the bathtub structure was also fitted in ‘flexsurv’ to the Lundy data.

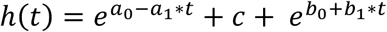

**Equation 1**. Hazard (mortality) function of the Gompertz bathtub model^30^. The first part of the equation (*a*_0_, *a*_1_, *c*) models the decline in mortality from early ages to adulthood. The increase in mortality with age is modelled by the second part of the equation, with *b*_0_ modelling the relative vulnerability to the increase in mortality (frailty) per *t*, as defined by the *b*_1_ parameter (ageing rate).

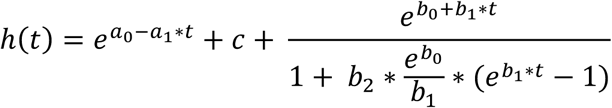

**Equation 2.** Hazard (mortality) function of the Logistic bathtub model^30^. The logistic model converges to the Gompertz model when parameter *b*_2_, which allows mortality to decelerate with age, approaches zero.

### Predictors of mortality in the Lundy population

#### Temperature data

We explored the relationships between the mortality of juveniles and resident adults with temperature. Minimum daily temperature data were obtained from the Meteorological Office (UK) for the weather station at RAF Chivenor (51.1° N, 4.1 W°, the closest official weather station to Lundy, 35 km to the east). We expected low temperatures to induce mortality. As a measure of how cold a year was, we used the number of days per census year (from April to March, as above) that had a minimum temperature below the lower daily minimum temperature quartile across the years 1999 to February 2012 (i.e. ≤ 4.6°C).

#### Presence of birds of prey

To assess a possible relationship between juvenile or adult mortality and the number of predators on the island, data on sightings of specified birds of prey were used to construct a predator index for each census year. Sightings of these species are routinely recorded by permanent island staff, members of the Lundy Field Society and visitors to the island, and are collated into monthly sighting numbers. We derived an annual index of all the sightings in each census year of sparrowhawk (*Accipter nisus*), merlin (*Falco columbarius*), hobby (*Falco subbuteo*) and kestrel (*Falco tinnunculus*) as the sum of the months in which there were at least two sightings of a species, divided by 48 (4 species times 12 months). This predator index therefore represents the relative proportion of a census year for which the population was at risk of predation. These raptor species commonly visit Lundy during migration, but in some years remain as residents, and their presence was therefore chosen as a proxy of the predation pressure acting on the population. All four species are known predators of house sparrows. In the case of the kestrel, which is generally known to have a preference for vole species, it is known that the species switches its diet towards passerines almost exclusively when the availability of small mammalian prey is reduced^35^. The only small mammal currently present on Lundy is the pygmy shrew (*Sorex minutus*), following the successful eradication of rats on the island in 2002–2004.

#### Statistical model

In order to separate the potentially shared covariance among the independent variables tested against mortality in the population, and to also correct for changes in age demography affecting adult mortality in a census year, linear mixed effects binomial models were fitted in ‘lme4’ in R. Two models were fitted, one for juvenile and another for adult mortality, that included a random intercept term for census year of the study and fixed terms for the three independent variables considered. For adult mortality, the effect of age on mortality was fitted as a factor, given the non-linear nature of this relationship (Figure 1), and individuals aged over 5 years were pooled into a single category to aid model convergence.

**Figure 1.**
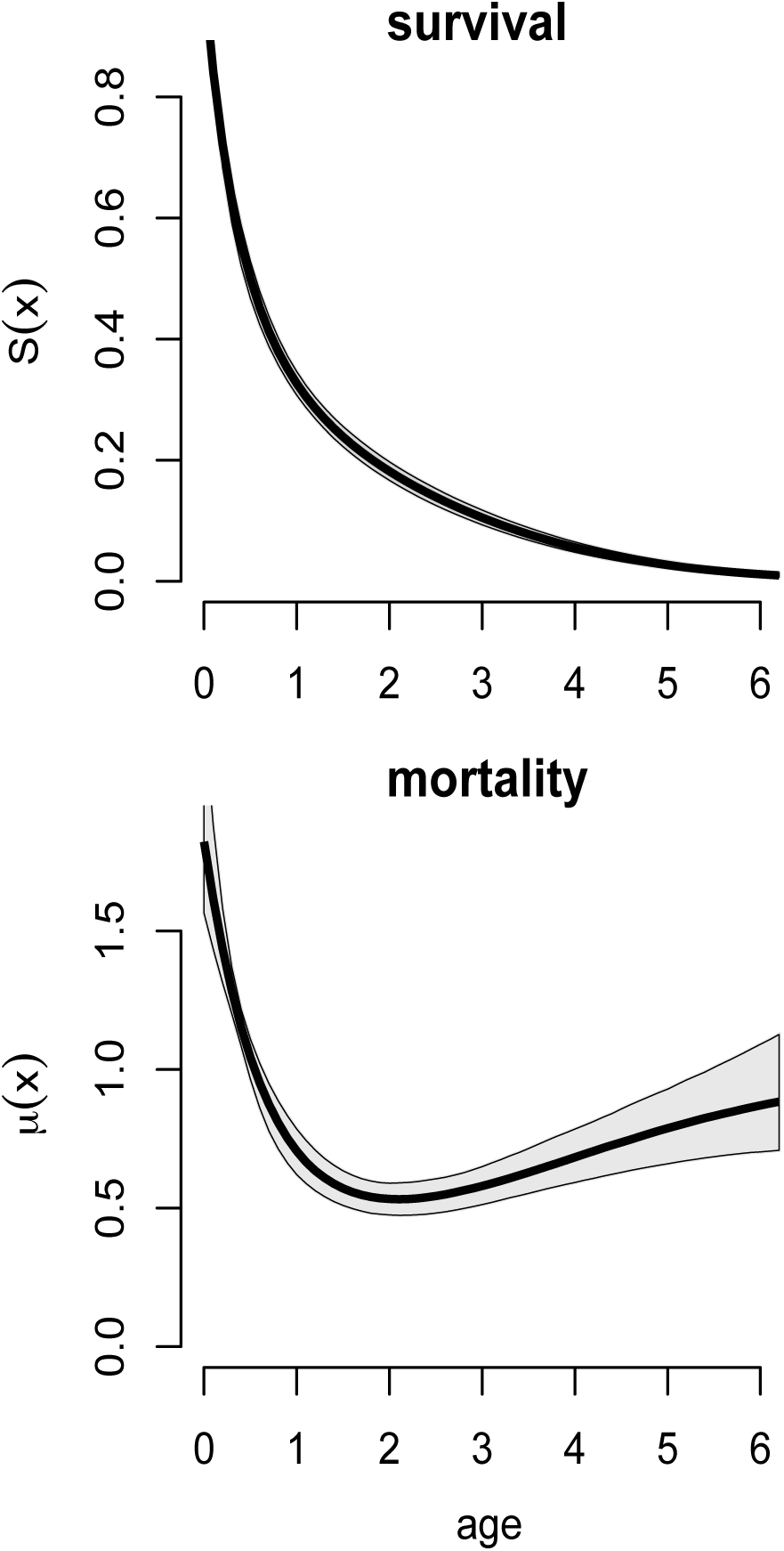
Fitted survival (top) and mortality (bottom) from the preferred model (Logistic, table 1) from day 12 (age 0) up to age 6, after which most individuals in the wild population have died. The grey shaded area indicates the 95% confidence interval around the estimate (solid line).

## Results

### Mortality trajectories and resighting probability

The Logistic model fitted the Lundy data best (Table 1, ΔDIC 32.9). The magnitude of the b2 parameter, with its 95% CI well above zero, indicated that mortality levelled off at the oldest ages, probably contributing to the superior fit of the Logistic model over the Gompertz model. There was no difference in the mortality trajectory between the sexes; Kullback–Leibner divergence calibrations remained very close to 0.5 for all parameters (range: 0.50–0.61). Mortality dropped steeply for adults that recruited into the population and there was a moderate acceleration in mortality with age, as indicated by the significantly positive values of the b1, the ageing rate parameter, in both the Logistic and Gompertz models (Table 1).

**Table 1.** Comparison of parametric survival models of the wild house sparrows on Lundy. The first three variables parameterise the bathtub shape (*a*_0_, *a*_1_ and *c*; Figure 1). The b set of adult mortality senescence parameters are defined accordingly (see text and Equation 1 and 2).

| Parameter | Gompertz (95% CI) | Logistic (95% CI) |
| --- | --- | --- |
| $a_0$ | 0.378 (0.150 : 0.640) | 0.407 (0.199 : 0.633) |
| $a_1$ | 1.923 (1.220 : 2.876) | 1.599 (0.944 : 2.530) |
| $c$ | 0.356 (0.095 : 0.526) | 0.285 (0.042 : 0.499) |
| $b_0$ ('frailty') | -2.690 (-4.368 : -1.322) | -3.224 (-4.778 : -1.828) |
| $b_1$ ('ageing rate') | 0.327 (0.154 : 0.548) | 0.749 (0.338 : 1.282) |
| $b_2$ ('deceleration') | | 1.045 (0.144 : 2.212) |
| DIC | 6537 | 6504 |

Estimated recapture probabilities were close to saturation and highly similar across all models (deviation of 0.001), with very narrow confidence intervals (for the preferred Logistic model = 0.96, 95% CI 0.95–0.97). Mortality in the captive population was lower and this was due to a change in the frailty parameter, with a highly similar rate of ageing in captivity and in the wild (Table 2, Figure 2).

**Table 2.** Gompertz fits of adult mortality in the captive and wild population. Note the high similarity in the ageing rate parameter (b_1_) and the substantial difference in the frailty parameter (b_0_). Refer to Figure 2 for a plot of the models.

| Gompertz Parameter | Captive (95% CI) | Wild (95% CI) |
| --- | --- | --- |
| $b_0$ ('frailty') | -3.117 (-3.461 : -2.769) | -1.309 (-1.423 : -1.194) |
| $b_1$ ('ageing rate') | 0.203 (0.127 : 0.270) | 0.228 (0.188 : 0.268) |

**Figure 2.**
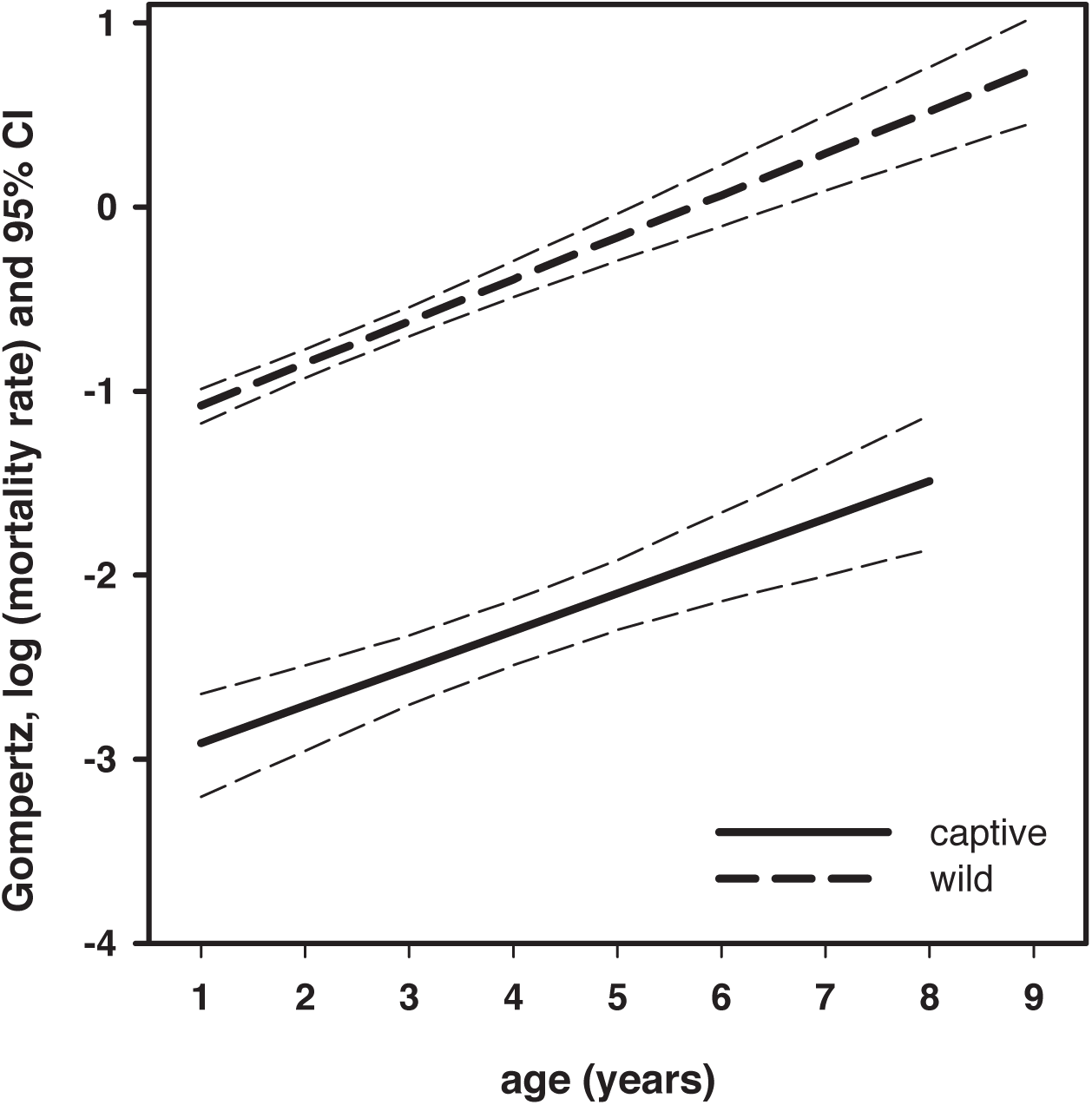
Fitted mortality rates under a Gompertz model of house sparrows in the wild and in captivity. Model predictions are plotted across the full range of ages available in the datasets.

### Detailed demography of juvenile and adult mortality

The high re-sighting probability and the near absence of dispersal to and from the island^24^ allowed us to separate juvenile mortality from adult mortality. Adults were assumed dead if not seen in the next year (ignoring the 4%, based on the estimated re-sighting probability, that we are expected to miss in each census year) and thus recruited juveniles could be separated from adult survival. Using this approach, we constructed a detailed picture of juvenile mortality and adult mortality independently, across the years of our study. Year to year variation in mortality (Figure 3) was high and statistically significant (logistic regressions, juvenile: χ^2^_(11)_= 49, p < 0.0001, adult: χ^2^_(11)_= 105, p < 0.0001). Note that some individual adults are used repeatedly in these analyses because they were alive in multiple years, creating pseudoreplication; yet, given the relatively short mean lifespan (Figure 1), we expect these effects to be relatively minimal. Juvenile and adult mortality covaried positively across years (*r*_*s*_ = 0.70, p = 0.015), indicating that when adult mortality was high, this was also the case for juveniles (Figure 3).

**Figure 3.**
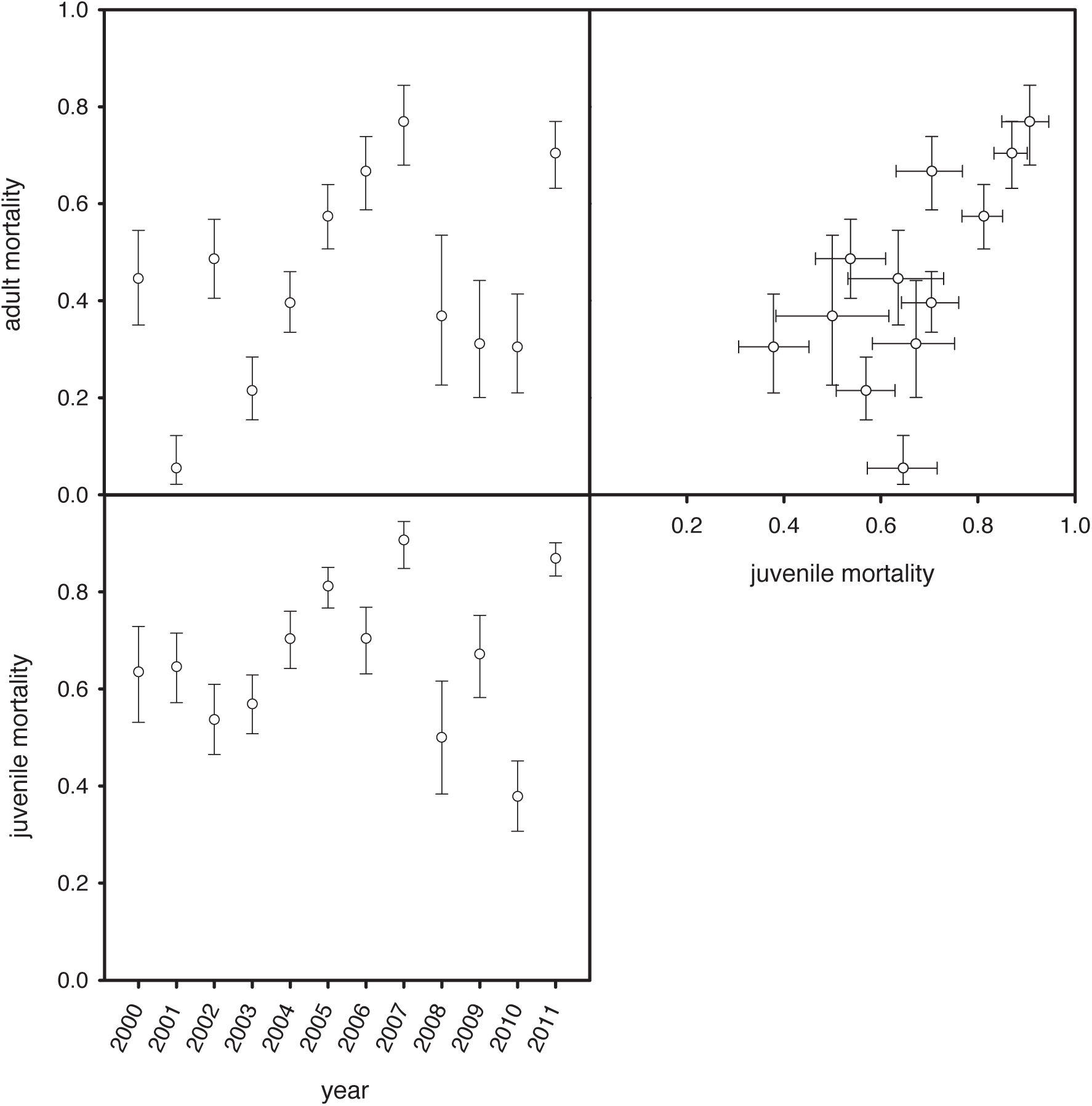
Variation in adult and juvenile mortality rates per census year (two graphs on left) and the correlation between these (top right graph). Whiskers indicate confidence intervals of proportions according to Blaker^57^, calculated using the package ‘PropCIs’ in R.

To explore any environmental effects on mortality, we investigated the effects of number of cold days per census year, population size, and predator index on adult mortality and juvenile mortality (Figure 4). Surprisingly, in cold years, adult and juvenile mortality was lower (*r*_*s*_ = −0.65, p = 0.02; *r*_*s*_ = −0.52, p = 0.08, respectively). Adult mortality and juvenile mortality were each positively related to population size (adult + juveniles), but neither relationship reached significance (adult: *r*_*s*_ = 0.27, p = 0.39; juvenile: *r*_*s*_ = 0.41, p = 0.19). Predator index predicted juvenile (*r*_*s*_ = 0.61, p = 0.03) and adult mortality (*r*_*s*_ = 0.59, p = 0.04). In the binomial mixed effect models used to pull apart the independent effects of these three independent variables, predator presence exhibited the strongest effects, with only this variable reaching significance and then only in adults (Table 3). This is probably due to the covariance between predator presence and cold weather (*r*_*s*_ = −0.38, p = 0.23), because separately each of these parameters did predict mortality significantly or showed a trend for both juvenile and adult mortality (Table 3).

**Table 3.** Estimates from the mixed binomial models, including census year as a random effect, either testing the three environmental variables together in a full model or separately, run for juvenile and adult mortality separately. Raw estimates of scaled variables are given with their standard errors. ** indicates *p* < 0.01, * indicates *p* < 0.05, † indicates *p* < 0.1. Models of adult mortality included age to correct for differences in age demography and its associated mortality between years.

|  | Juvenile mortality |  | Adult mortality |  |
| --- | --- | --- | --- | --- |
|  | full model | separate | full model | separate |
| **cold days** | $-0.23 \pm 0.19$ | $-0.41 \pm 0.18^*$ | $-0.12 \pm 0.24$ | $-0.41 \pm 0.25^\dagger$ |
| **population size** | $0.19 \pm 0.17$ | $0.35 \pm 0.19^\dagger$ | $0.13 \pm 0.22$ | $0.30 \pm 0.27$ |
| **predator index** | $0.21 \pm 0.18$ | $0.34 \pm 0.18^\dagger$ | $0.56 \pm 0.24^*$ | $0.65 \pm 0.23^{**}$ |

**Figure 4.**
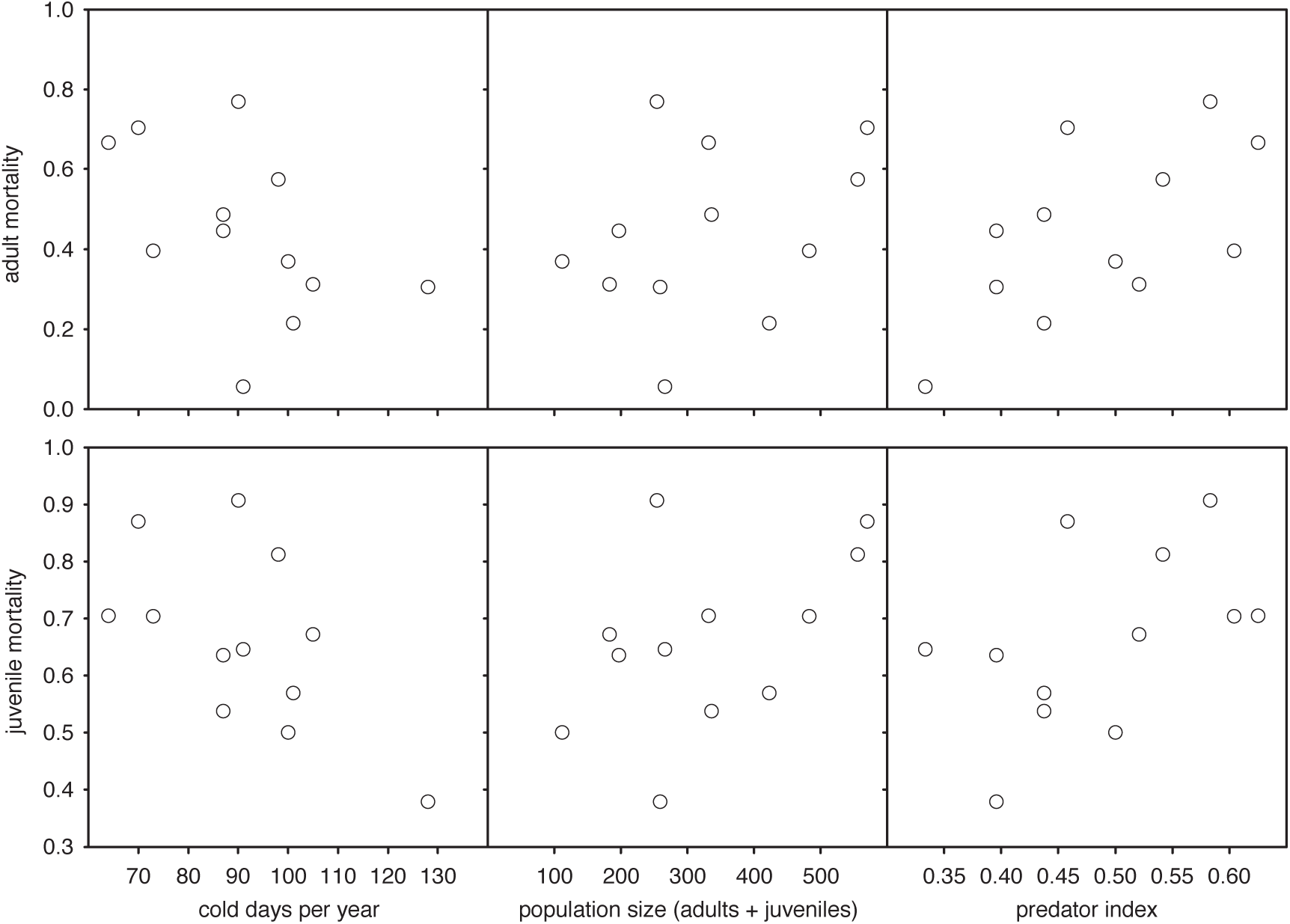
Scatter plots between adult or juvenile mortality and temperature, population size or the presence of raptors. Significant relationships were present between predation pressure and both juvenile and adult mortality, and mortality of both juveniles and adults was higher in warmer winters. See text for rank correlations of these relationships and Table 3 for binomial mixed models.

## Discussion

### Actuarial senescence

We detect relatively small but significant mortality senescence (‘actuarial senescence’) in wild house sparrows (Figure 1). There was a ∼1.6-fold increase in mortality from its trough at age ∼2 to age ∼6; in comparison, there is a ∼3-fold increase in mortality rates in a US human population from the age of 60 to 100 years and a ∼2.7-fold increase in mortality rate in male mice from the age of 1.4 to 2.9 years^36^. This level of actuarial senescence in the sparrow population is detected despite strong environmental effects on adult and juvenile mortality. This is relevant to the evolution of senescence, given that the strength (and sometimes direction) of selection on life history is changed by the level of stochastic (environmental) noise^37,38^. Extrinsic mortality shapes optimal investment in the soma over reproduction (current versus future reproduction trade-off)^39^ and, accordingly, different levels of extrinsic mortality on the population level lead to different levels of senescence-related mortality^14^. When this selective pressure – extrinsic mortality – is more variable, selection on intrinsic, senescence-related, mortality is weaker and hence a larger standing variation in intrinsic mortality is expected^40^. Moreover, different levels of stochasticity can also lead to the evolution of differential bet-hedging strategies^41^.

Studies of the fitness costs of senescence^7^ and evolutionary theory of ageing^42^ have not considered environmental stochasticity. Different levels of environmental stochasticity might also explain differences between species in the demography of fertility and mortality^12,43^. Understanding the effects of environmental variation in extrinsic mortality on intrinsic mortality and/or their interactions will be a next step in understanding the evolution of senescence in the wild. In addition, such considerations might explain why the rates of reproductive senescence and mortality senescence do not always match^7,14,44,45^, perhaps because the selective pressures maintaining them are differentially susceptible to environmental effects. Effects of the developmental environment on reproductive, but not survival senescence^13^ is perhaps an illustration of such differential environmental effects on life-history traits.

### Environmental effects on the demography of mortality

House sparrows in captivity and in the wild show a highly similar rate of ageing but differential frailty, consistent with similar comparisons in mammals, namely baboons^18^ and humans^20^. This suggests that the ageing rate is a specific property of a species, insensitive to environmental effects. An invariable within-species rate of ageing is fundamental to the compensation law of mortality and the mortality deceleration that follows from the reliability theory of ageing^16,46,47^. Although comparative evidence from birds (this study) and mammals^18,20^, including the mortality deceleration observed in this study and others^4^, now point in this direction, in contrast, experimental evidence suggests that ageing rate can be flexible. Dietary restriction in rodents reduces the rate of ageing without affecting frailty^16^, contrary to, for example, the effect of dietary restriction on mortality in *Drosophila melanogaster*, which is exclusively attributable to a change in frailty^48,49^. Parental effort also modulates the ageing rate in the wild, as demonstrated using brood size manipulations in jackdaws (*Corvus monedula*)^11^, the white-throated sparrow (*Zonotrichia albicollis*)^50^, and the Seychelles Warbler (*Acrocephalus sechellensis*)^51^. This suggests that, to some extent, the wide variation in mortality trajectories between species^12^ can be due to differential environmental or population effects^52^. Distinguishing between captive and wild populations^53,54^, and examining experimental effects on frailty and ageing rate independently, will be crucial steps towards understanding the evolution of mortality trajectories and senescence.

### Juvenile and adult mortality

The high recapture probability we estimated from the BaSTA models allowed for the separation of juvenile and adult mortality, which in many other study systems is not possible. Juvenile and adult mortality were also found to be sensitive to environmental effects, and covaried positively, suggesting that juveniles and adults died of similar environmental causes. The analysis also suggests that any density dependent effect is relatively limited, given that adult survival does not impinge on juvenile mortality. Therefore, food availability and aggression, resulting from competition for food or mating territories, are unlikely to limit the population size.

Indeed, population size in a census year did not impinge strongly on either juvenile or adult mortality (Table 3, Figure 4). Of the other two environmental variables, predator presence was most predictive of mortality, especially in adults. When considered separately, cold weather was associated with improved survival of juveniles and adults. Irrespective of whether this is indirect via predation pressure, it goes against the usual expectation of harsher winters, although still relatively mild on Lundy, causing more mortality. Without being able to attribute causes of death to starvation, disease or directly to predation, we can only speculate on the roles of physiology and/or ecology in the relationship between mortality and ambient temperature.

Intensive monitoring, as in the Lundy house sparrow population, achieving near perfect re-sighting rates, is key when inferring biology from the demography of mortality, which is crucial in explaining the evolution^12^ and biology^3,55,56^ of ageing. The confounding factors of dispersal and unmarked individuals that are typical of most study populations would bias such estimates, and can lead to potentially false conclusions about differences in demography between species when the degree of these biases differs across study populations and species^12^. Understanding the physiology and evolution of relatively invariable species-specific ageing rates in the face of strong environmental effects, as we show here for a bird in addition to existing evidence from mammals, is pivotal to understanding natural selection on senescence and its physiology.

## Acknowledgements

We thank Ian Cleasby, Duncan Gillespie, Simon Griffith, Yu-Hsun Hsu, Maria Karlsson, Maria-Elena Mannarelli, Nancy Ockendon and Claire Prosser for their contributions to the fieldwork. We are grateful to Bart Kempenaers for providing access to the captive population of house sparrows and for support. We thank Emmi Schlicht and Silke Laucht from the Kempenaers Group for providing mortality data for the population, and the animal caretakers for looking after the birds. We thank the UK Meteorological Office for supplying the daily temperature data and the Lundy Company staff for their enthusiastic support of our research. The Lundy Field society and visitors to the island are thanked for recording and collating sightings of birds on Lundy. Support by the UK Natural Environment Research Council (NE/J024597/1 and NE/M005941/1 to TB). SN was supported by a Rutherford Discovery Fellowship (New Zealand) and a Future Fellowship (Australia). MJPS is supported by a Sir Henry Wellcome Fellowship and a Sheffield Vice Chancellor’s fellowship. TB was supported by a Leverhulme Fellowship.

